# *ILIAD*: A suite of automated Snakemake workflows for processing genomic data for downstream applications

**DOI:** 10.1101/2023.10.11.561910

**Authors:** Noah Herrick, Susan Walsh

## Abstract

**Background:** Processing raw genomic data for downstream applications such as imputation, association studies, and modeling requires numerous third-party bioinformatics software tools. It is highly time-consuming and resource-intensive with computational demands and storage limitations that pose significant challenges that increase cost. The use of software tools independent of one another, in a disjointed stepwise fashion, increases the difficulty and sets forth higher error rates because of fragmented job executions in alignment, variant calling, and/or build conversion complications. As sequencing data availability grows, the ability of biologists to process it using stable, automated, and reproducible workflows is paramount as it significantly reduces the time to generate clean and reliable data.

**Results:** The *Iliad* suite of genomic data workflows was developed to provide users with seamless file transitions from raw genomic data to a quality-controlled variant call format (VCF) file for downstream applications. *Iliad* benefits from the efficiency of the Snakemake best practices framework coupled with Singularity and Docker containers for repeatability, portability, and ease of installation. This feat is accomplished from the onset with download acquisitions of any raw data type (FASTQ, CRAM, IDAT) straight through to the generation of a clean merged data file that can combine any user-preferred datasets using robust programs such as BWA, Samtools, and BCFtools. Users can customize and direct their workflow with one straightforward configuration file. *Iliad* is compatible with Linux, MacOS, and Windows platforms and scalable from a local machine to a high-performance computing cluster.

**Conclusion:** *Iliad* offers automated workflows with optimized time and resource management that are comparable to other workflows available but generates analysis-ready VCF files from the most common datatypes using a single command. The storage footprint challenge of genomic data is overcome by utilizing temporary intermediate files before the final VCF is generated. This file is ready for use in imputation, genome-wide association study (GWAS) pipelines, high-throughput population genetics studies, select gene candidate studies, and more. *Iliad* was developed to be portable, compatible, scalable, robust, and repeatable with a simplistic setup, so biologists who are less familiar with programming can manage their own big data with this open-source suite of workflows.

## Background

It is estimated that genomics research will produce approximately 40 exabytes of data within this decade alone [1]. Genomic data processing is paramount to deciphering functional information, particularly through the identification of candidate trait and disease-associated genetic variants within population data that are highly relevant in clinical, forensic, and other biological fields of research. Genome-wide association (GWA) and candidate gene studies in particular continue to pave the way toward our understanding of human health and disease [2, 3]. These computational studies require genotypic information derived from a number of platforms and data types such as microarray and sequence data, but the files obtained from these platforms require significant computational resources to perform format conversions before analyses can even take place. For example, when a researcher receives single nucleotide polymorphism (SNP) array data generated from a commercial platform (i.e., Illumina) each sample’s BeadArray data is stored in the form of green and red IDAT files. The decryption of the stored summary intensities for every probe type on an array must be performed using proprietary software, either the Illumina Array Analysis Platform Genotyping Command Line Interface (IAAP-CLI) or GenomeStudio programs (Illumina, Inc. San Diego, CA, USA) which can be labor intensive, as processing is limited to the number of samples in a single sample sheet [4]. Each program introduces steps that may impede researchers without a computational background from working with their own big data.

Next-generation sequencing is responsible for another source of genomic data that is on a larger scale. Sequence data, whether open source or provisional access, has a foray of different file types including raw sequence FASTQ files and highly compressed alignment (CRAM) files. These files are unreadable using standard text editors and software and therefore require several computationally intensive steps including alignment, variant calling, ID tagging, and sample/dataset file consolidation before investigators can include them in their analyses. At present, there are several standardized reference population datasets available. A few of the most popular include 1000 Genomes Project [5], Human Genome Diversity Project [6] (HGDP), and Simon Genome Diversity Project [7]. Although this sequence data is open source, it requires hours of processing before use, leading to delays that can impact research. Lower costs in wet laboratory sequencing consumables and SNP arrays have also opened the door towards in-house data preparation for researchers interested in using sample sequence information as both a study target and reference panel resource [8]. This is further supported by the use of imputation software servers such as the Michigan Imputation Server [9], Sanger Imputation Server [10], and more data privacy permitting imputation pipelines such as *Odyssey* [11] to generate even larger amounts of genetic information than what was originally genotyped. This data shift signals a focal point that researchers need to overcome, which is the lengthy and arduous task of manual genomic data handling. For example, a singular sample with paired-end reads approximately 50 gigabytes (GB) in combined size can lead to manual processing times of up to 48 hours. It can be an extremely daunting task for the biologist who prepared the sample for sequencing to later process the genetic output, particularly when it may be thousands of samples.

Several workflows and pipelines have been developed over the years to simplify this task by integrating third-party bioinformatics software tools and reducing runtimes. These workflows, however, will often be catered for one data file type [12–14]. A recently developed workflow called OVarFlow [15] is based on the Snakemake [16] common workflow language for processing variant data from high-throughput sequencing FASTQ files. It was designed to be customizable and can be used to analyze a wide range of variant data, including single nucleotide variants (SNVs), indels, and structural variants (SVs). The pipeline includes a number of different steps under GATK [17] software for processing and analyzing variant data, including quality control, alignment, variant calling, annotation, and filtering. There are several manual steps that must be performed such as downloading the FASTQ data and reference genome files and updating the Snakemake version before the workflow can start. Although it is a valuable resource for processing FASTQ input data, it would benefit greatly from additional modules to accommodate the other data file types that biologists typically encounter such as CRAM, IDAT, and VCF.

Variant call format (VCF) files are the standard output of processed sequence data and the prerequisite to many genetic data analysis tool kits. As there is a rise in user-generated data and collections of populations that are available in varying data formats online, we have the potential to unite these datasets to develop more globally represented custom reference datasets. This is an important emerging capability for researchers that are well-equipped with quick and easy data processing pipelines. It allows them to better represent select populations with improved imputation accuracies and therefore advance big data genome analyses on a global scale. Improvements such as increased imputation accuracy have already been noted using these approaches and methodologies [18]. It is essential therefore to have versatile workflows that can handle as many raw genomic data formats as possible, from start to finish including an ability to merge datasets from different genome assembly builds, whether sequence or SNP array generated. The capability to clean and efficiently combine datasets into singular genetic files in an automated fashion will make research between groups easier, ultimately increasing power in the detection of variants in larger association studies. There are workflow languages, such as Snakemake, that greatly reduce manual processing steps by automating them into a pipeline. It is no longer practical in genomics for big data processing to rely on segmented data handling scripts and manual handling of excessive intermediate files. Workflow management systems are highly valuable for this automation.

Here we introduce *Iliad*, a suite of Snakemake workflows developed with several modules for automatic and reliable processing of raw or stored genomic data that lead to the output of ready-to-use genotypic information necessary to drive downstream applications. *Iliad* offers a containerized workflow with optional automatic downloads of desired files from file transfer protocol (FTP) sites coupled with the use of any genome reference assembly for variant calling using BCFtools [19]. All dependencies are pre-installed or systematically downloaded and/or built when invoked to considerably reduce the time and effort required to execute the workflows. At present we demonstrate usage comparisons with another genetic data processing workflow and show time-saving improvements as well as increased data input flexibility using human sequence data only, but also provide instructions on how this pipeline can be adapted to cater for other genomes. *Iliad*’s minor startup requirements and complementary modules for quality control and data cleaning support its user-friendly and customizable characteristics. With compatibility for the major operating systems in mind, *Iliad* offers a scalable solution from local machines to high-performance computing (HPCs) clusters to address the needs of any genomics researcher.

## Implementation

### Pipeline architecture and configuration file

Genomic data processing poses a challenge for genetic research studies because it involves multiple program dependency installations, vast numbers of samples with raw data from various Next Generation Sequencing (NGS) platforms, and inconsistent genetic variant ID and/or positions among datasets. The *Iliad* suite of genomic data workflows automates the central steps in genomic data processing for several NGS data types with implementation through the Singularity [20] container system and Snakemake workflow management system. These systems form the basis of *Iliad* and account for its ease of distribution, reproducibility, and scalability to ultimately accommodate users with a simplified and standardized suite of workflows that are easy to implement.

All but one of the required program dependencies for *Iliad* are contained within a pre-built singularity image file available on Sylabs cloud (library://ncherric/iliad/igdp-container:latest) that is automatically pulled down into the workflow by Snakemake. A Docker container solution is also provided (https://hub.docker.com/repository/docker/ncherric/iliad/general) with options for AMD64 or ARM64 architectures. The Illumina IAAP-CLI is not permitted for distribution, so the download link is accessible to *Iliad* and users via the configuration file. The Singularity definition file and Dockerfile used to build the containers are also provided with *Iliad* on Github and ReadTheDocs in case there are any user-specific modifications to build a custom version. The container base system is Ubuntu 20.04 [21] Linux and includes BWA [22] for read mapping, SAMtools [19] and PicardTools [23] for user-choice of sorting and compressing the alignment files, BCFtools for variant calling, +gtc2vcf BCFtools plug-in [24] for converting SNP array files, and miniconda [25] for creating rule-based conda environments as needed. The latest *Iliad* container contains up-to-date versions of SAMtools and BCFtools, however, users can download previous versions if preferred by using a tag that corresponds to the version of software (e.g library://ncherric/iliad/igdp-container:v1.14).

*Iliad* functions under Snakemake best practices and takes advantage of several useful features. *Iliad* works as an inference pipeline where the user can specify the desired endpoint, and Snakemake will then infer which rules are to be run based on the following user input: the final invocation of the Snakemake command, existing input/output files in the workflow, and a primary configuration file. The desired endpoint is declared as the input to the “rule all” found in each of the independent workflows’ main Snakefile. This is already pre-set based on the goal of each workflow. There are certain Snakemake flags that will affect the starting point, such as ‘--forceall’ which triggers all rules to be executed whether there are existing files or not. In a scenario where *Iliad* has been invoked and progress was interrupted, the workflow will begin where it left off saving both time and resources. This offers users a flexible entry point into the workflow, meaning researchers can begin using *Iliad* even if they possess data files corresponding to input that is mid-stream of a workflow.

All *Iliad* workflows refer to the primary configuration ‘config.yaml’ file which has numerous controller variables with clearly denoted purposes. This file is the central point of customizing user needs. Many of the adjustments are binary conditions that are dependent on files a user may already possess. The most important configurations can be found at the top of this primary file. For example, adding the working directory path location of the *Iliad* directory is required. Once that single change is made in the configuration file, *Iliad* will be able to execute a demo based on tutorial data. This provides users with an instant glimpse into the workflow and how it works, making it easier to begin working with new data. A thorough how-to guide for each workflow and a demonstration of the tool is provided on “Read the Docs” (https://iliad.readthedocs.io/en/latest/).

### Modularization

*Iliad* was developed as a suite of workflows using the modularization capabilities of Snakemake (Figure 1). It includes data-specific pipeline modules that are designed for raw sequence data (FASTQ), Illumina SNP array data (IDAT), and a common storage format for sequence alignment data (CRAM). Each of these data types are common sequence data files. The user is free to choose which of these data processing modules best suit their needs by executing the corresponding Snakefile(s). Each of these modules provide flexible start and end points depending on any pre-existing data files and the Snakemake flags included in the command line invocation. Additionally, *Iliad* features independent submodules for lifting over reference assembly genomic positions (GRCh37 to GRCh38 and vice versa) and merging multiple VCF files at once. These submodules can be used independently or combined within the raw sequence processing modules.

**Figure 1.**
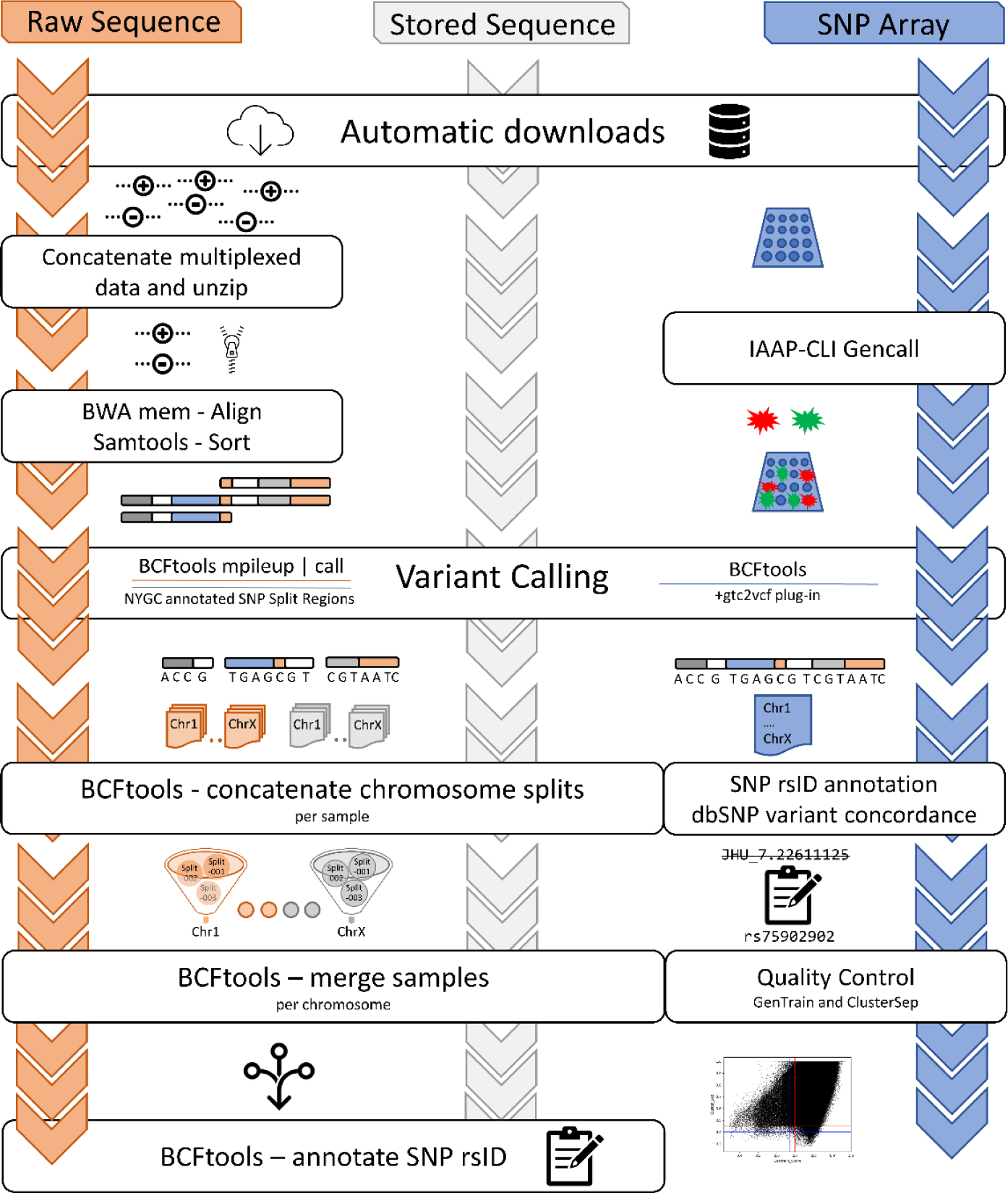
Workflow schematic for each of the modules. A user can run the modules independently or simultaneously. The sequence data modules, raw FASTQ and stored CRAM, follow the same base set of rules after variant calling.

### Computational specifications

Development and benchmarking took place on Carbonate [26] and Ulysses, respectively. Carbonate is Indiana University’s large-memory computer cluster designed for data-intensive tasks. It is home to a cluster of 72 Lenovo NeXtScale nx360 M5 server compute nodes each with 12-core Intel Xeon E5–2680 v3 CPUs and 256 GB of RAM. There are additionally 8 large-memory compute nodes containing 512 GB of RAM, but those were not necessary for the purpose of *Iliad*. Ulysses is a remote server with more immediate access for the Department of Biology at Indiana University-Purdue University Indianapolis (IUPUI). Ulysses is comprised of two 16-core Intel Xeon Gold CPUs. Both Carbonate and Ulysses surpass the hardware necessary to run *Iliad.* It is also possible to execute *Iliad* workflows on a local machine such as a desktop or laptop equipped with Linux, MacOS, or Windows operating systems (OS) possessing at least 1 core, 16 GB of RAM, and enough disk space storage to sustain the amount of data needed. We recommend having as much computational storage as possible for sample processing, but this will vary depending on the nature of the research. We provide the minimum system requirements necessary to execute the demo tutorial. RAM and CPU usage metrics were collected using the collectl utility (http://collectl.sourceforge.net/) and the built-in ‘benchmark’ declarative in Snakemake.

### Raw Sequence Read Data Workflow

The raw sequence read data module is comprised of 24 rules (Table S1; Figure S1), not including the rule ‘all’, which is designated as the driver of the workflow to provide the final desired output VCF. Each of these rules has been optimized using the “resource” directive, allotting for a specific time and memory request specific to each rule. Some rules may be branched into hundreds of jobs based on the number of samples. Job scheduling systems are more likely to run a queued job with smaller dedicated time and memory requests for smaller tasks. Some of the rules, such as downloading annotations files, are only necessary to be run once. *Iliad* will re-use those general annotation files for later runs and cache them so that the other *Iliad* modules have access and can skip redundant downloads. There are instances when input data, such as a reference genome assembly FASTA file, is already locally available on a system. If so, adding ‘true’ to the ‘IhaveReference’ variable in the configuration file will trigger its automatic use. Similarly with input FASTQ reads or CRAM alignment files, placing the data into the appropriate directory and declaring the ‘--ignore-incomplete’ flag in command line invocation is all that is required if a user does not need the automatic downloading feature.

The main workflow handles raw sequence data which can be used for reference or target data. The first advantage is a download checkpoint that uses ‘wget’ to acquire the user’s FASTQ data from the specified URL in the config file. Since many studies include multiplexed sequencing runs across many lanes, all the files associated with a particular sample name will be downloaded into a temporary ‘downloads’ folder where they will be accessed for a concatenation rule and output as one set of unzipped paired-end reads into the ‘fastq’ folder. If the user already has FASTQ data placed in the ‘fastq’ directory, the checkpoint will be satisfied. Quality control of the FASTQ data is performed via FASTQC [27], and reports are generated in HTML format which is a valuable step for users to check the raw sequence quality of downloaded data. *Iliad* will proceed with completing other rules based on the input data. Since read mapping is next for FASTQ data, the rule for obtaining the reference genome indicated in the configuration file will begin to produce the remaining and necessary input files that the read mapping rule requires. The reference genome is retrieved using a script modified from the reference wrapper, ‘0.74.0/bio/reference/’, in the “dna-seq-gatk-variant-calling” workflow found on the Snakemake workflow catalog [28]. With a specific reference assembly and corresponding index file, *Iliad* will then begin read mapping using the burrows-wheeler alignment (BWA) package (v0.7.17). The main workflow, then, pipes the BWA output (SAM file) to Samtools ‘sort’ and creates a sorted BAM file. Sorting of the file through a pipe eliminates the need for a higher storage capacity by reducing intermediate files. After this step, the main workflow implements the BCFtools variant caller to produce VCFs that contain the genotypic information. Due to user configuration included in this workflow, variant calling parameters set for BCFtools can be adjusted if the user wishes to set specific thresholds or flags for the ‘mpileup’ and ‘call’ algorithms. The same is true if application of the ‘norm’ flag is desired. This allows *Iliad* to keep up to date with any new developments in BCFtools variant calling scripts.

Rather than performing variant discovery and calling all possible variants found in the alignment file, *Iliad* utilizes a curated list of variants (n = 120,046,375) from the New York Genome Center [29] to perform genotyping as this list comprises of stable variants observed across the 1000 Genomes Project dataset. A rule in the main workflow automatically downloads these files from the associated FTP site [30]. There is one file per chromosome, 23 in total. The next rule splits each chromosome into equally divided chromosomal regions to further mitigate the computational reading and writing time observed when using BCFtools ‘view’ on one chromosome file in its entirety. The number of chunked region files can be customized in the configuration file by the user to fit any system-specific requirements. Any ambiguity in chromosome naming conventions is also handled within this rule. The chunking methodology is used to drastically increase speed either in series or in parallel, however, using workers in parallel is the faster approach. We chose BCFtools as our variant calling software because it provides a wide array of filtering options and useful plugins that maximize user customization and data flexibility. Within the *Iliad* workflow this gives users the ability to modify BCFtools commands as needed, particularly when new versions of the software become available. It has consistently been one of the preferred performance-evaluated variant calling tools [31–34] for sequencing data whilst including a capability for analyzing other data types (i.e., microarray data). Furthermore, the implementation of BCFtools concatenation and merging features complement the chunking methodology to optimize VCF generation of multiple samples from whole genome sequence (WGS) data.

### Stored Sequence Read Data Workflow

Sequence alignment files (SAM or BAM), especially for the human genome, create impending hard disk storage challenges that can become quite costly. Therefore, compressed sequence alignment (CRAM) files are a very popular storage file format commonly found on publicly available project FTP sites such as 1000 Genomes Project [5], Human Genome Diversity Project [6], and Simons Genome Diversity Project [7]. The CRAM file format is continually undergoing modifications and updates to improve speed and accuracy [35] therefore this workflow is particularly up to date with developments in data compression. *Iliad* incorporates a stored sequence read data module that downloads desired open-source CRAM data from a server and performs the above-mentioned steps for variant calling on the retrieved files, just as it would perform variant calling on sorted BAM files in the raw sequence module. This is a critical module that supports in-house development of WGS reference panels and enables a fast and efficient addition of standard reference data sets that are publicly available. *Iliad* is one of the first Snakemake workflows that specifically manages the automation of CRAM to VCF data processing using BCFtools and user-controlled software flags. It is important to note, especially for new users, the exact same genome reference assembly that was used by the research group that generated the CRAM data is required. *Iliad*’s configuration file provides a binary variable to declare which reference genome assembly must be used and the reference genome file path if it must be supplied by the user. For example, variant calling CRAM files from HGDP [6] require the ‘GRCh38_full_analysis_set_plus_decoy_hla.fa’ reference genome.

### SNP Array Data Workflow

An important feature of this pipeline is the ability to integrate the processing of SNP array data from its raw IDAT form. Typically for Illumina-specific SNP array files, the data must either be uploaded to a Windows only software (GenomeStudio) or utilize numerous command line applications. To simplify this entire process and seamlessly integrate Illumina SNP array data processing into our workflows, we containerized the open-source programs and included download steps for programs with end user license agreements, such as IAAP-CLI.

With the proper tools in the workflow environment, the procedure to obtain a VCF is facilitated by passing the raw data through the appropriate conversion steps. Initially, the data must be physically located in the “./Iliad/data/snp_array/idat/” directory, so that the IDAT files can be converted to GTC files using IAAP-CLI ‘gencall’. There are product and support files that assist this conversion which include manifest and cluster files. It is imperative that the user knows which reference assembly their study will need and to indicate as such in the configuration file for their automatic retrieval. The Infinium Multi-Ethnic Global-8 v1.0 microarray web links for product and support files corresponding to *Homo sapiens* GRCh37 and GRCh38 reference assemblies are included and may provide assistance to users locating their specific array support documentation. The correct product and support files will be used as input for follow-up third party tools to continue the conversion.

A BCFtools plugin, ‘+gtc2vcf’ is included in the singularity container and is called to convert all sample GTC files into one VCF file located in the same directory (“./Iliad/data/snp_array/gtc/”). Illumina support files, acquired automatically, update obscure loci names to widely accepted rsIDs. Although this may be a sufficient endpoint for some analyses, we include a filtering step that will find overlapping rsIDs from the dbSNP annotation VCF from NCBI [36]. Users can configure which dbSNP file corresponds with your desired genome reference assembly to produce a quality VCF that can be easily joined with other genomic data using standard rsIDs. Finally, raw IDAT files provide metadata for GenTrain and ClusterSep scores for every variant and can be used to filter out calls of poor quality. At present, there are default upper and lower thresholds for GenTrain (0.7 and 0.67) and ClusterSep (0.45 and 0.4) built into the workflow, but these can be adjusted according to user preference. Final outputs include a summary graph and a quality controlled VCF file.

### Submodules for additional VCF optimization

To best serve additional data processing applications, *Iliad* features several submodules that assist with build conversions and the merging of VCF files with other datasets available to the user e.g., reference data. Variant genomic positions largely depend on the reference assembly used for alignment, therefore datasets from different sources may have varying VCF ‘POS’ fields that inevitably represent the same SNP but cannot be merged correctly. The most comprehensive submodule functions as an automatic Lift-over and Merge task (Figure 2). Users simply “drag and drop” their datasets into the directory (‘./Iliad/data/vcf_Lift-and-Merge’), provide a project name and reference assembly preference in the configuration file, and list the files to merge in the text file (‘./Iliad/config/mergeTheseVCFs.txt’). Doing so results in a tidy project space dedicated to a specific project and build conversion. The submodule enables users to perform multi-VCF merges on compressed or decompressed data represented by *Homo sapiens* GRCh37 or GRCh38 positions. Once configuration variables have been set and the ‘Lift-and-Merge_Snakefile’ executed, the pipeline detects and updates genomic positions, naming conventions, and initiates the merge of autosomes and the X chromosome using BCFtools. The submodule also automatically detects which version (GRCh37 or GRCh38) each VCF file is before passing it to the correct processing channel for generation of the final build expressed by the user. Processes to filter the independent VCFs and quality check the final merged VCF then occurs, also based on user configuration (maximum SNP and individual missingness).

**Figure 2.**
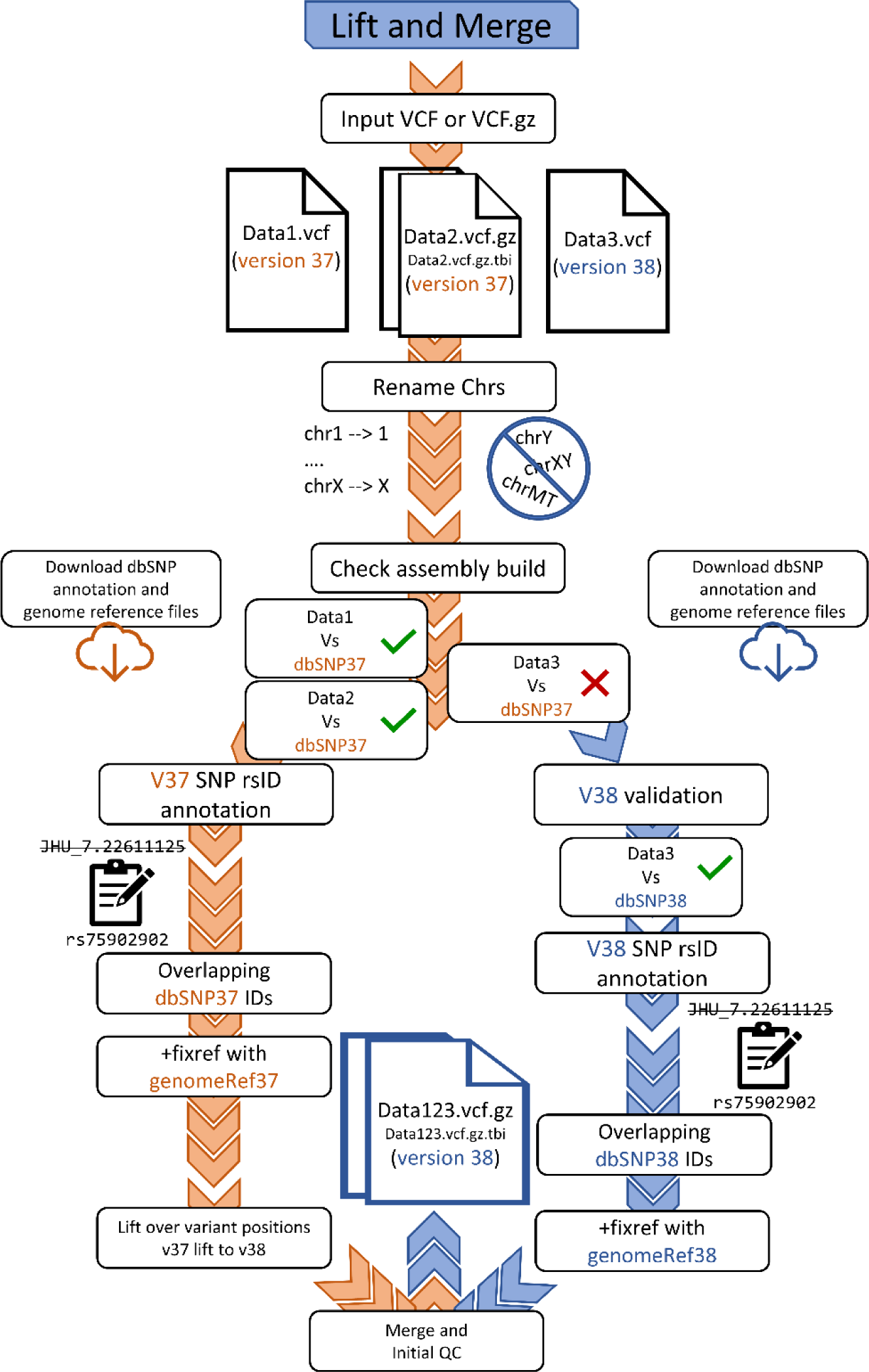
Workflow schematic for the Lift and Merge submodule. VCF data files from independent datasets with genomic positions that reflect either the *Homo sapiens* GRCh37 or GRCh38 genome reference assemblies can be merged. This workflow is specifically for a final merged VCF file configured to have genomic positions in GRCh38. The orange denoted pathway represents GRCh37 data input and the blue denoted pathway represents GRCh38 data input.

The Lift-over and Merge options also have split submodules for ease of use. There are two independent lift-over options that convert *Homo sapiens* GRCh37 positions to *Homo sapiens* GRCh38 positions and vice versa. The ‘liftoverTo37_Snakefile’ or ‘liftoverTo38_Snakefile’ must be passed to your Snakemake command and VCF files migrated to the ‘./Iliad/data/liftover’ directory. These perform a lift-over on VCF(s) but do not merge them. The data merging submodule can be performed on its own or with integration in the main sequence and SNP array modules with the execution of the ‘targetRefMerge_Snakefile’. This is especially useful when processing both reference and target data for a particular analysis.

## Results and Discussion

*Iliad* was designed to simplify the arduous task of downloading and converting raw sequence data from thousands of individuals for both WGS and SNP array data into a single optimized and clean VCF for downstream applications using an all-in-one suite of workflows customizable for the user. A greater in-depth how-to guide hosted by “Read the Docs” (https://iliad.readthedocs.io/en/latest/) makes the process extremely simplified for biologists who may not be as comfortable working with the large datasets they may generate. Baseline estimates of time, computational resources, and storage required for the three main *Iliad* modules are provided in Tables S1-3 (see Additional File 1). Additional testing occurred on a number of platforms including Google Cloud Platform (GCP), Windows, and MacOS. Summarized evaluation metrics (Table 1) illustrate the resource and time estimates a user will need. One must be prepared with ample disk space when working with big data genomics, however, *Iliad* mitigates storage challenges by eliminating unnecessary intermediate files. After running through the tutorial data in the raw sequence module, roughly 33 GB is stored. Approximately 28 GB includes the reference genome assembly and annotation files, while the remaining 5 GB is the resulting VCF data footprint for the tutorial data of a single sample (paired-end reads; FASTQ). Tests with 1 to 5 CRAM samples from 1000 Genomes Project resulted in a 14.3 to 21.6 GB VCF file. An example run of the SNP array module generated a VCF of 17.2 GB for 190 in-house samples typed using the Illumina MEGA array (n = 1,686,450 SNPs). These 190 in-house samples are not provided in our demo due to Institutional Review Board (IRB) restrictions. Ultimately, *Iliad*’s novelty is established as an all-in-one suite that is managed by a single configuration file with the ability of the workflow to flexibly commence at any phase of data processing due to the Snakemake framework. Many annotation files will be reused by other processes, so they are conveniently cached and available to any and all of the workflows within *Iliad,* including the submodules.

**Table 1.** Computational resources and time necessary to perform each of the main workflows with the described sample data. These benchmarks were recorded from the Ulysses HPC and run without a job scheduler. *Provided tutorial data as an auto-download for demonstration purposes when *Iliad* is retrieved.

| Workflow | Initial file type | Tutorial sample | Tutorial sample size (GB) | Maximum RAM (GB) | Disk space (GB) | Cores | Time (hours) |
| --- | --- | --- | --- | --- | --- | --- | --- |
| Raw Sequence (~12x) | FASTQ | KPGP-00127* | 17.3 | 10 | 33 | 32 | 7.6 |
| Stored Sequence (~30x) | CRAM | NA12718* | 14.3 | 0.5 | 34 | 32 | 37.3 |
| SNP Array | IDAT | 190 In-house samples | 10.6 | 3 | 45.2 | 16 | 3.4 |

With regards optimal memory and timesaving performance as well as storage outcomes, a comparison of *Iliad*’s Raw Sequence module and OVarFlow [15] was also conducted (Figure 3). It is important to note that OVarFlow’s main purpose is variant discovery which would require additional processing and filtering steps over *Iliad’s* selective variant genotyping, and this was the closest workflow comparison available to record the time elapsed while processing raw sequence reads to obtain a clean, ID-annotated VCF file in an automated fashion. For OVarFlow’s installation and ease of use, the Conda environment required a Snakemake version update from 5.26.1 to 7.8.5 in order for it to begin the run. It also required the user to manually download the reference genome; a GFF file - if not present with the reference genome, and input FASTQ data. These extra manual preparation steps required more user intervention than *Iliad,* typically adding at least 60 minutes of computational task time.

**Figure 3.**
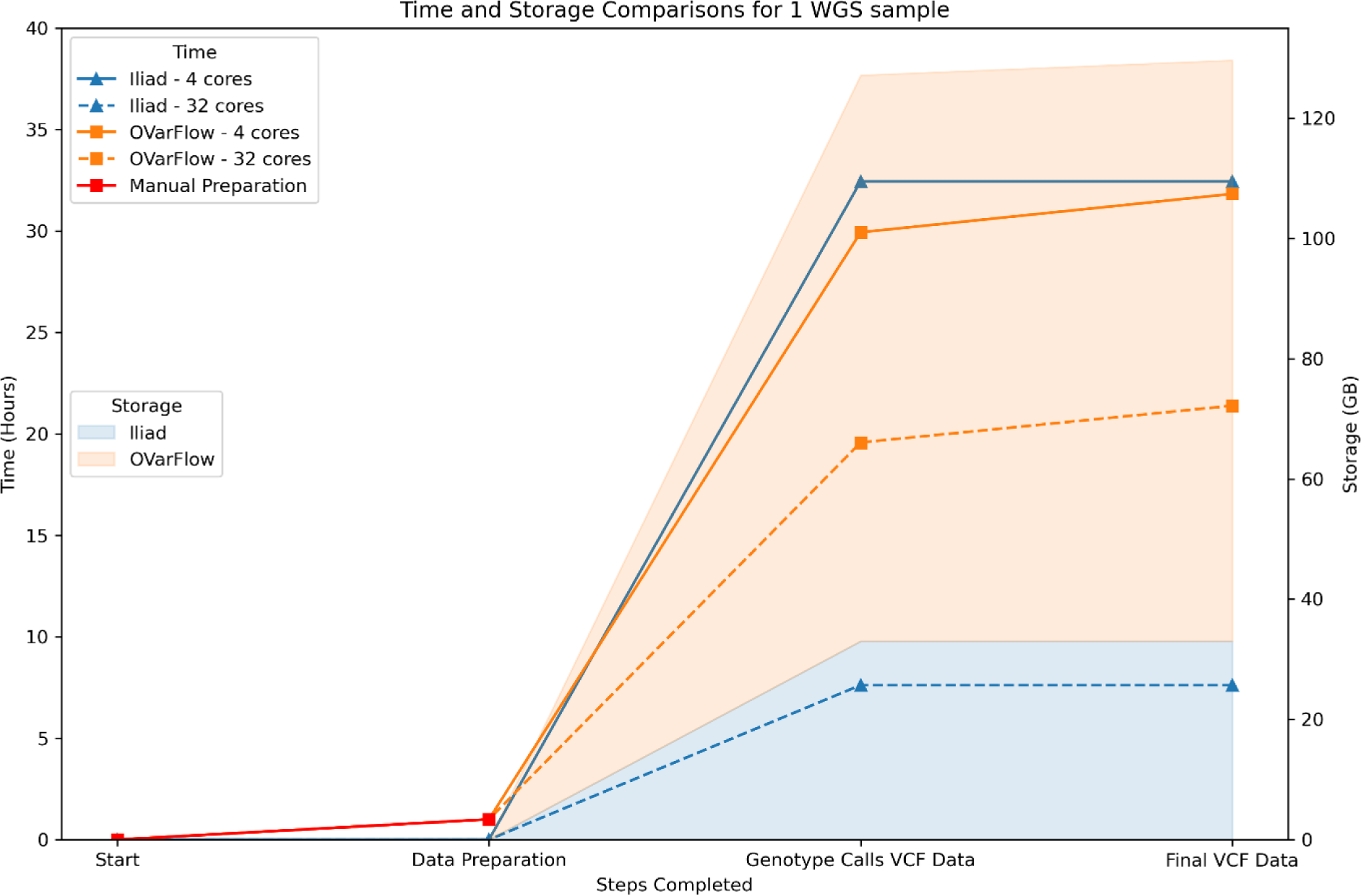
Time and storage comparisons between *Iliad*’s Raw Sequence Read Data module and OVarFlow. The left Y-axis represents the time elapsed in hours for the selected steps completed (X-axis) by each of the workflows at 4 and 32 cores. The right Y-axis and shaded area of the graph represent the amount of cumulative storage in gigabytes (GB) for the selected steps completed (X-axis) by each of the workflows, regardless of the number of cores. The Genotype Calls VCF Data and the Final VCF Data are the same for *Iliad*’s Raw Sequence Read Data workflow.

Overall, the two pipelines were completed within 0.5 hours of each other for paired-end reads of one sample from the open-source Korea Personal Genome Project [37, 38] (KPGP) using 4 cores for 4 jobs in parallel. Within this figure, we also provide information on the time it took for OVarFlow to generate a combined VCF for all variant calls (labeled ‘Genotype Calls VCF Data’) before pipeline completion, to show OVarFlows performance prior to unique variant filtering. This gave a difference of 1.9 hours between both pipelines.

However, *Iliad* quickly gained time improvements with increasing processing power and at 32 cores it finished nearly 3 times faster for 32 jobs in parallel at 7.62 hours compared to OVarFlow’s 21.4 hours. This is due in part to the default processing power settings of OVarFlow that is set to 6 cores and *Iliad* which is set to 12 cores. *Iliad* capitalizes on more cores because of the flexibility in adding more chromosome splits for variant calling using BCFtools. Several other factors may also contribute to these time differences such as the subtle differences in variant calling between BCFtools and GATK. Worthy to note, however, is the difference in storage observed after pipeline completion. *Iliad* contains many clean-up steps that are intrinsic to the workflow that reduce the storage footprint for raw sequence data processing by a factor of 4 when compared to OVarFlow, which can aid users if they are processing hundreds of samples at once.

The time expenditures for *Iliad* when job sizes were scaled to 5 and 10 samples were recorded at 17.5 hours and 31 hours, respectively, when using 32 cores for 32 jobs in parallel. Time and resource usage of the chromosome splitting methodology was recorded across multiple combinations of splits and cores allocated (Figure S4). Chromosome 22 from sample KPGP-00127 from the open-source KPGP repository was used for testing. The limiting factor of *Iliad*’s speed was simply the supplied number of cores. Significant advantages lie in the reuse of commonly used files, such as annotation files and genome reference assemblies. For instance, rerunning *Iliad* to retrieve and process new samples amounted to decreased run times in comparison to a clean install simply due to file download limitations from the human genome servers [39], thus reaffirming its efficiency. Although we were unable to directly compare our other main modules as we could not find any other comparable pipeline that focuses on CRAM files and IDAT files, we do offer benchmark results in Tables S2 and S3 for future comparisons.

In sum, although time assessments are quite comparable given the number of variant calls generated in both VCFs using WGS data (Table 2), the main difference between the pipelines is in their utility for downstream applications. *Iliad* performs variant calling on specific genomic positions detailed in a region file in an effort to combine and clean datasets of multiple builds, cohorts and datatypes, whereas OVarFlow utilizes genotyping for more stringent variant discovery on a single dataset. *Iliad* has been designed to facilitate data generation for downstream GWAS and candidate association studies that require large numbers of individuals for increased power. It offers a general easy-to-use genomic data processing workflow that provides human genetic researchers greater accessibility to a set of variants across the genome as opposed to variant discovery. However, if researchers choose and wish to alter BCFtools commands (as provided in this pipeline), they may opt to exclude region files and perform exhaustive variant calling on the entire available sequence information, thus facilitating the capture of additional alternative variant calls found in the datasets.

**Table 2.** Comparison of genotype and variant data type between *Iliad* and OvarFlow calculated with BCFtools ‘stats’ flag. The ‘Genotype Calls VCF’ represents the count of records in Step “06_combined_calls” within OVarFlow. The ‘Final VCF’ (records) generated from OVarFlow’s pipeline is the reduced variant-only file. *Iliad* produces a singular final VCF within its pipeline that calls a specific set of annotated genomic positions from NYGC [30]. A breakdown of the number of SNPs and INDELs within the VCF is reflective of the pipelines’ respective variant callers.

| Workflow | Genotype<br>Calls VCF | Final VCF | SNPs | INDELs |
| --- | --- | --- | --- | --- |
| *Iliad*'s Raw Sequence Module | - | 119,960,362 | 9,520,169 | 498,893 |
| OVarFlow | 610,405,020 | 4,034,429 | 3,623,864 | 411,214 |

*Iliad* submodule workflows may prove the most useful to researchers with multiple VCF files that normally require meticulous data wrangling prior to merging. For biologists who do not have the time to conduct repetitive tasks and troubleshoot small data discrepancies on multiple datasets, the submodules built into *Iliad* are of extreme value due to their simplistic and timesaving properties. File aggregation across all data types and projects using a singular pipeline is a prime example of this. Users can simply ‘drop and run’ by putting several VCF files into one folder to merge them into a single VCF regardless of compression, genome build, ‘CHR’ naming conventions, file size, and number of SNPs or samples. This capability is available for all researchers but may be of particular interest for those that want to merge tens to hundreds of data files (VCF) that they may have at their disposal; a time-consuming and confusing task is now made very easy and efficient. The final steps in the merger also include quality control measures which are specified by the user and sum up an extremely beneficial and time efficient module. Demo data [40–42] was not provided with *Iliad* for this submodule, although, example data can be freely obtained from the Estonian Biocentre (https://evolbio.ut.ee/). Example datafiles from published literature [40–42] were downloaded in Bed, Bim, and Fam Plink [43, 44] formats from this site and converted into VCF files using Plink2 [43, 44]. Benchmarks are provided using this demo data (Table 3) to give users an insight into the resources required, and replication if needed. Manual dataset compilation is a demanding process whether the data consists of a small number of genotypes or WGS information. The Demo data consisted of genome-wide microarray data (n = 1,286,187 SNPs). The ability to generate a single quality controlled VCF from multiple files is an attractive workflow on its own, but combined with its ability to automatically detect which build and how to process the files greatly enhances the scope of *Iliad’s* suite of workflows.

**Table 3.**
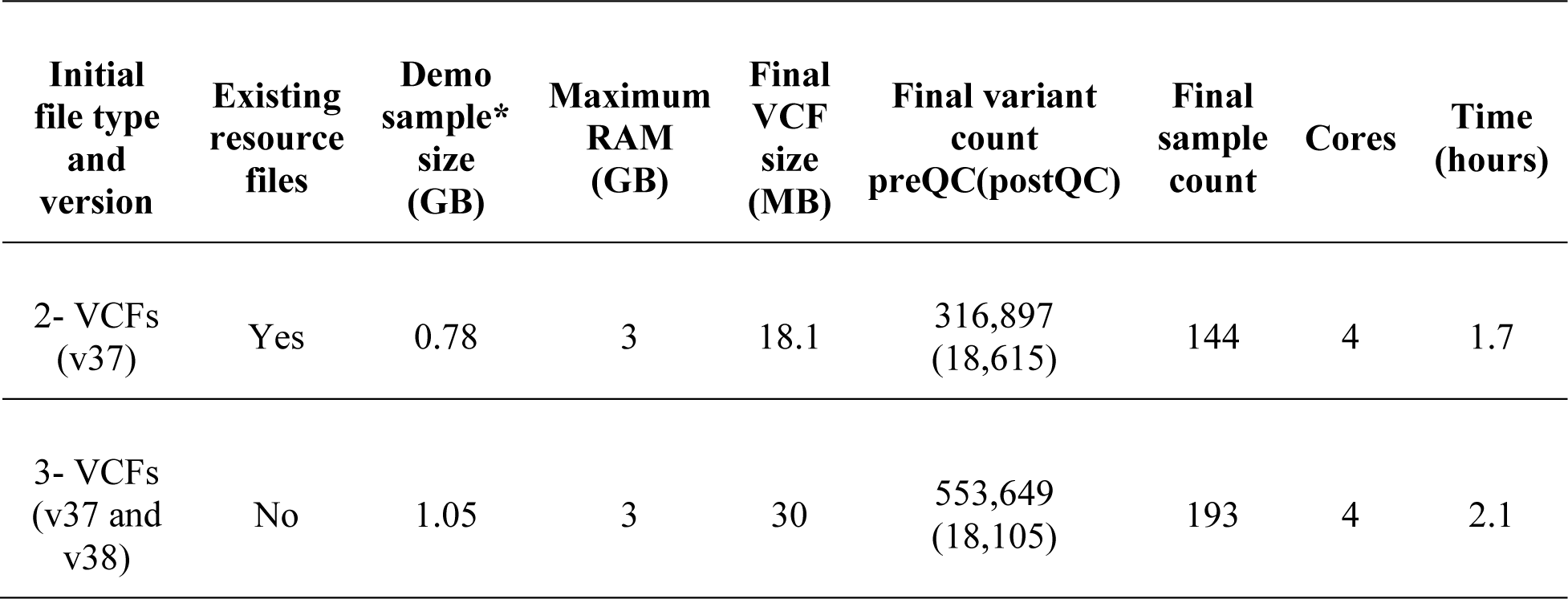
Computational resources and time necessary to perform the lift over and merge submodule on demonstration data [40–42] without any genome build information given. The data can be accessed from the Estonian Biocentre (https://evolbio.ut.ee/) or from data availability instructions associated with each study. The default Quality Control (QC) was set to remove variants and individuals with > 5% missingness.

As for the future development of *Iliad*, it will be expanded to include other data file types such as Affymetrix’s CEL SNP array file types (Affymetrix, Inc. Santa Clara, CA, USA) in addition to the inclusion of other variant callers such as GATK or FreeBayes [45] should the user have specific needs. Additionally, human genomic data was used in the development, testing, and benchmarking of this study, however, it will also be possible to process genomic data from other model organisms using *Iliad* if the necessary genome reference was specified and certain features adapted. Future versions of *Iliad* aim to accommodate users with these enhancements.

## Conclusion

The need for genomic data processing is expected to vastly increase based on continual NGS cost efficacy. Variant data files are a standardized solution for genotypic information derived from raw sequence data in a controlled and reduced format, but its generation is complicated by numerous software installations and program versions, disjointed file formats, and a lack of workflow consistency among researchers. *Iliad* standardizes this process by containerizing the required software tools and streamlining the entire workflow, whilst also leaving room for user quality control preferences. Accompanied by visual outputs of raw data quality, *Iliad* is the first workflow suite of its kind to simplify and automate the management of genomic data processing that will highly benefit biologists newer to the bioinformatics field.

## Availability and Implementation

The *Iliad* genomic data pipeline is open source and can be found on GitHub (https://www.github.com/ncherric/Iliad). It is easy and straightforward to setup on several operating systems using containers and is considered an ‘out-of-the-box’ suite of workflows thanks to the thorough documentation and visual how-to guides that complement *Iliad* (https://iliad.readthedocs.io/). Program dependencies and external downloads of supplementary files are automatically facilitated by *Iliad*. This suite of genomic data processing pipelines was tested using Windows, MacOS, and HPC Linux systems using both Singularity and Docker containers.

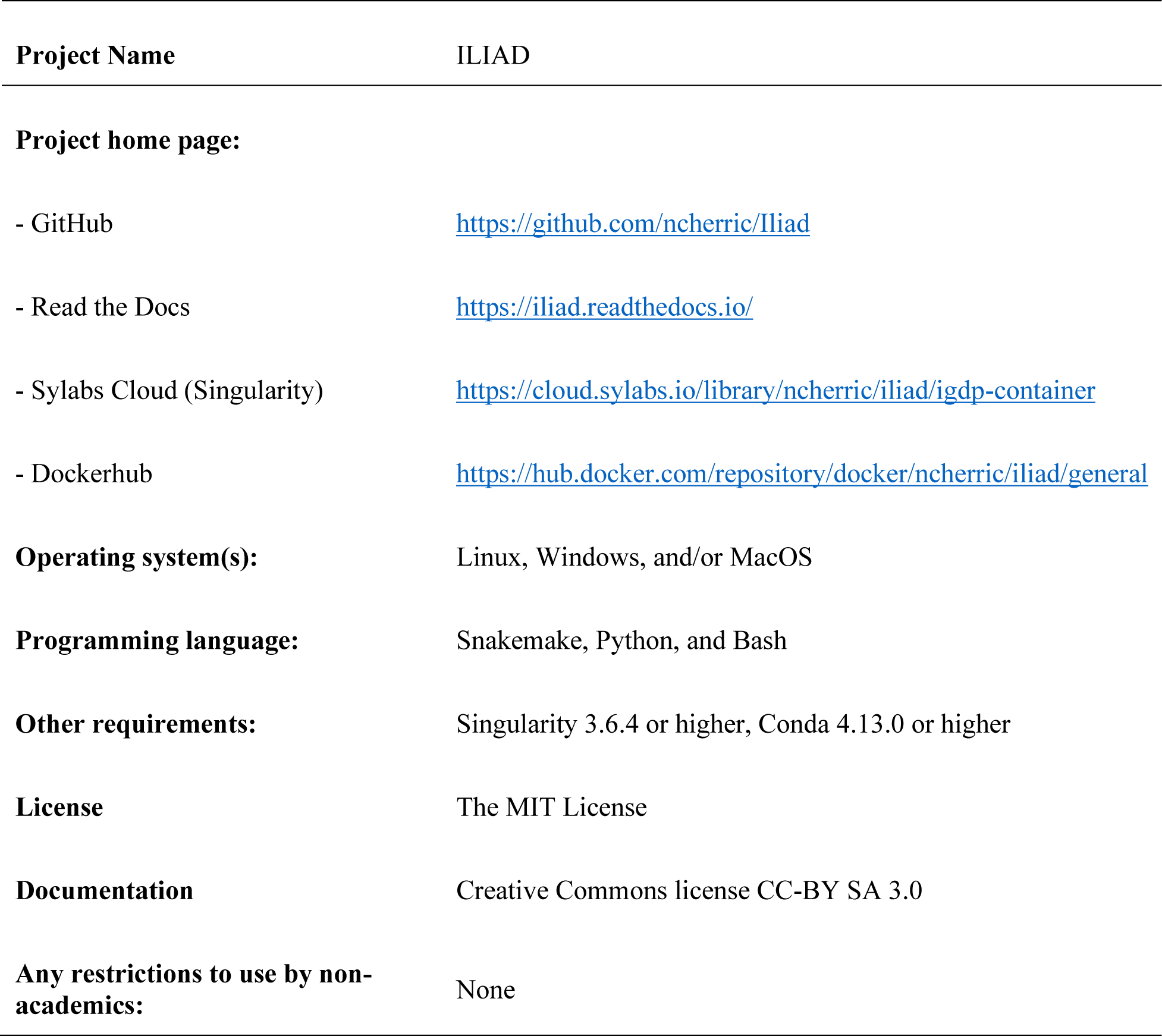

|  |  |
| --- | --- |
| **Project Name** | ILIAD |
| **Project home page:** |  |
| - GitHub | https://github.com/ncherric/Iliad |
| - Read the Docs | https://iliad.readthedocs.io/ |
| - Sylabs Cloud (Singularity) | https://cloud.sylabs.io/library/ncherric/iliad/igdp-container |
| - Dockerhub | https://hub.docker.com/repository/docker/ncherric/iliad/general |
| **Operating system(s):** | Linux, Windows, and/or MacOS |
| **Programming language:** | Snakemake, Python, and Bash |
| **Other requirements:** | Singularity 3.6.4 or higher, Conda 4.13.0 or higher |
| **License** | The MIT License |
| **Documentation** | Creative Commons license CC-BY SA 3.0 |
| **Any restrictions to use by non-academics:** | None |

## List of abbreviations

GWAS: Genome-wide association study
SNP: Single nucleotide polymorphism
IAAP-CLI: Illumina Array Analysis Platform Genotyping Command Line Interface
CRAM: Compressed Reference-oriented Alignment Map
HGDP: Human Genome Diversity Project
GB: gigabyte
SNV: Single nucleotide variant
SV: Structural variant
VCF: Variant call format
FTP: File transfer protocol
HPC: high performance clusters
NGS: Next-generation sequencing
rsID: Reference SNP cluster identifier
FASTA: Sequence data file format
FASTQ: extension of FASTA to include sequence quality data along with sequence data
IDAT: Intensity data
IUPUI: Indiana University-Purdue University Indianapolis
BAM: Binary alignment map; Sequence alignment map
WGS: Whole genome sequence
BWA: Burrows-Wheeler aligner
IRB: Institutional Review Board

## Declarations

### Ethics approval and consent to participate

Not applicable.

### Consent for publication

Not applicable.

### Availability of data and materials

Whole-genome sequence data used in development and testing was derived from the following open-source databases: The 1000 Genomes Project [5, 29] and The Korean Personal Genome Project [37, 38]. Genome-wide Array data (IDAT) used in development and testing was collected and is protected under Indiana University IRB Protocol 1409306349. Demo data files [40–42] used in the Lift and Merge submodule were obtained from the Estonian Biocentre (https://evolbio.ut.ee/) open-source data repository.

### Electronic supplementary material

Additional_File_1.pdf

## Authors’ contributions

NH designed and programmed the *Iliad* pipeline, wrote the guidance literature on how to run *Iliad*, and drafted the manuscript. SW oversaw the project and helped draft the manuscript.

## Acknowledgments

We would like to give special thanks to all the open-source data repositories and study volunteers for making this research possible and to Frankie Wilke, Chad Pressler, Kyra Mullins, Ryan Eller, and Rob Hart for testing various modules of the *Iliad* suite of Snakemake workflows.

