## Additional File 1 for "*ILIAD*: A suite of automated Snakemake workflows for processing genomic data for downstream applications"

### Supplementary Material

Herrick, N. Walsh, S.

**Table S1.** Time and resource usage for *Iliad*'s raw sequence read data module. Each rule in the Snakemake workflow utilized the "benchmark" directive to collect this data. Some rules will constitute a number of jobs depending on the wildcards e.g. chromosomes and number of splits per chromosome. For this reason, values for time and resource usage were averaged across the jobs per rule. The benchmarks were conducted with paired-end reads of one sample, KPGP-00127, from the open-source Korea Personal Genome Project (KPGP: http://kpgp.kr) on a dedicated remote server in the Department of Biology at Indiana University – Purdue University Indianapolis. This run was conducted using 8 cores provided to Snakemake for 8 jobs in parallel. No internal rule parallelization occurred; however, it is possible and greatly speeds up some rules e.g. alignment. Since the "benchmark" directive is rule-specific for resource monitoring, there was no impact on any independent rule's time and resource report from performing these benchmarks with 8 jobs in parallel.

| Rule | Jobs (with 5 splits) | Average values across Jobs per Rule | | | | | | | | |
|---|---|---|---|---|---|---|---|---|---|---|
| | | h:m:s | max_rss | max_vms | max_uss | max_pss | io_in | io_out | mean_load | cpu_time |
| alignment | 1 | 2:02:25 | 19083.97 | 20901.21 | 19071.88 | 19075.59 | 12.76 | 42668.80 | 1186.09 | 232.43 |
| annotate_rsIDs_with_dbSNP | 23 | 0:20:44 | 70.60 | 395.50 | 61.68 | 64.75 | 11.64 | 54.34 | 98.54 | 1226.49 |
| annotation_index | 23 | 0:00:02 | 3.36 | 6.99 | 3.44 | 3.44 | 2.63 | 0.00 | 62.19 | 1.99 |
| bwa_index | 1 | 0:51:34 | 4477.62 | 4486.36 | 4469.03 | 4471.97 | 10.88 | 6651.68 | 99.17 | 3069.47 |
| cat_splits_to_chr_list | 23 | < 0:00:01 | 0 | 0 | 0 | 0 | 0 | 0 | 0 | 0 |
| check_for_chr_string | 1 | 0:00:01 | 24.41 | 28.84 | 21.45 | 23.00 | 8.98 | 0 | 0 | 0.42 |
| concat_reads | 1 | 0:05:51 | 4.37 | 14.71 | 2.28 | 3.37 | 2.64 | 49940.04 | 88.06 | 167.87 |
| concat_splits_per_chrom | 23 | 0:00:19 | 1462.44 | 7012.67 | 1452.97 | 1456.09 | 11.66 | 448.54 | 87.93 | 10.68 |
| concat_splits_per_chrom_index | 23 | 0:00:02 | 3.31 | 6.95 | 3.41 | 3.41 | 2.40 | 0 | 57.14 | 1.88 |
| download_NYGC_annotations_file | 23 | 1:23:16 | 4.31 | 9.35 | 4.38 | 4.38 | 3.17 | 3092.38 | 1.76 | 89.07 |
| download_dbSNP_file | 1 | 0:07:16 | 8.79 | 13.42 | 6.49 | 7.95 | 4.23 | 15607.06 | 33.24 | 0.09 |
| fnPath_to_file | 115 | < 0:00:01 | 0 | 0 | 0 | 0 | 0 | 0 | 0 | 0 |
| get_genome | 1 | 0:04:44 | 53.21 | 1362.86 | 41.79 | 45.18 | 16.06 | 2380.64 | 13.57 | 38.46 |
| merge_sample_chrom_list | 23 | < 0:00:01 | 23.15 | 418.16 | 20.82 | 21.89 | 8.42 | 0 | 0 | 0.39 |
| merge_samples_per_chrom | 23 | 0:00:25 | 44.25 | 223.59 | 35.42 | 38.49 | 11.50 | 36.50 | 67.21 | 15.09 |
| merge_samples_per_chrom_index | 23 | 0:00:02 | 3.33 | 6.98 | 3.42 | 3.42 | 2.52 | 0 | 60.35 | 1.93 |
| split_chroms | 115 | < 0:00:01 | 0.51 | 2.62 | 1.54 | 1.54 | 2.03 | 30.22 | 0 | 0.15 |
| split_chroms_chr_string | 115 | < 0:00:01 | 0.55 | 2.62 | 1.56 | 1.56 | 2.04 | 69.42 | 0 | 0.25 |
| tabbed_regions | 23 | 0:00:02 | 41.75 | 450.92 | 33.03 | 36.49 | 9.38 | 42.65 | 103.49 | 2.94 |
| tabbed_regions_chr_string | 23 | 0:00:02 | 40.72 | 339.76 | 32.91 | 36.03 | 10.71 | 64.81 | 67.59 | 2.74 |
| unique_regions_file | 23 | 0:00:11 | 4.70 | 10.25 | 2.37 | 3.79 | 1.98 | 67.80 | 99.26 | 11.62 |
| variant_calling | 115 | 0:21:30 | 231.96 | 347.63 | 218.21 | 222.68 | 11.80 | 10.83 | 99.51 | 1289.78 |
| variant_calling_index | 115 | < 0:00:01 | 3.19 | 36.91 | 3.38 | 3.38 | 2.67 | 0.00 | 3.05 | 0.43 |
| wget_reads_from_url | 1 | 0:16:59 | 4.39 | 9.35 | 4.41 | 4.41 | 3.17 | 17565.03 | 18.27 | 186.41 |

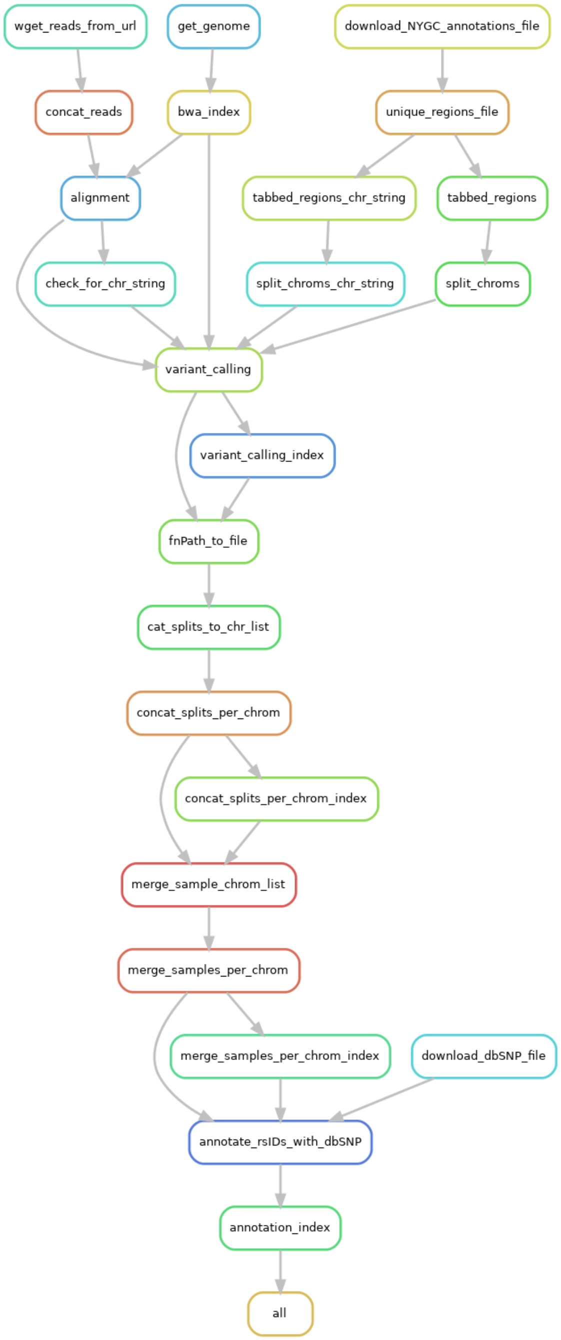


**Figure S1**. A directed acyclic graph for Iliad's raw sequence read data module.

**Table S2.** Time and resource usage for *Iliad*'s stored sequence read data module. Each rule in the Snakemake workflow utilized the "benchmark" directive to collect this data. Some rules will constitute a number of jobs depending on the wildcards e.g. chromosomes and number of splits per chromosome. For this reason, values for time and resource usage were averaged across the jobs per rule. The benchmarks were conducted on one sample, NA12718, in CRAM format from the 1000 Genomes Project (ftp://ftp.sra.ebi.ac.uk/vol1/run/ERR323/ERR3239480) on a dedicated remote server in the Department of Biology at Indiana University – Purdue University Indianapolis. This run was conducted using 16 cores provided to Snakemake for 16 jobs in parallel. No internal rule parallelization occurred. Since the "benchmark" directive is rule-specific for resource monitoring, there was no impact on any independent rule's time and resource report from performing these benchmarks with 16 jobs in parallel.

| Rule | Jobs (with 5 splits) | Average values across Jobs per Rule | | | | | | | | |
|---|---|---|---|---|---|---|---|---|---|---|
| | | h:m:s | max_rss | max_vms | max_uss | max_pss | io_in | io_out | mean_load | cpu_time |
| annotate_rsIDs_with_dbSNP | 23 | 0:31:40 | 73.66 | 666.09 | 64.91 | 67.97 | 103.95 | 68.71 | 99.18 | 1885.39 |
| annotation_index | 23 | 0:00:04 | 2.39 | 4.98 | 2.47 | 2.47 | 2.14 | 0.01 | 25.75 | 2.66 |
| bwa_index | 1 | - | - | - | - | - | - | - | - | - |
| cat_splits_to_chr_list | 23 | < 0:00:01 | 11.84 | 846.32 | 10.90 | 10.91 | 0.00 | 0.00 | 0.00 | 0.01 |
| check_for_chr_string | 1 | 0:00:01 | 36.50 | 1434.71 | 33.45 | 34.97 | 21.24 | 0.10 | 35.07 | 0.80 |
| concat_splits_per_chrom | 23 | 0:00:35 | 1533.85 | 6999.38 | 1524.93 | 1528.00 | 23.09 | 366.06 | 78.72 | 16.36 |
| concat_splits_per_chrom_index | 23 | 0:00:03 | 12.43 | 793.84 | 11.77 | 11.78 | 2.97 | 0.00 | 72.20 | 2.96 |
| cram_variant_calling | 115 | 5:46:43 | 585.14 | 2038.99 | 568.49 | 573.08 | 136.39 | 120.02 | 99.86 | 16570.40 |
| variant_calling_index | 115 | 0:00:01 | 8.84 | 440.94 | 8.48 | 8.48 | 3.19 | 0.02 | 40.11 | 0.58 |
| download_NYGC_annotations_file | 23 | - | - | - | - | - | - | - | - | - |
| download_dbSNP_file | 1 | - | - | - | - | - | - | - | - | - |
| fnPath_to_file | 115 | < 0:00:01 | 23.92 | 1548.80 | 22.14 | 22.16 | 0.00 | 0.00 | 0.00 | 0.03 |
| get_genome | 1 | - | - | - | - | - | - | - | - | - |
| merge_sample_chrom_list | 23 | 0:00:01 | 23.24 | 660.61 | 20.77 | 21.73 | 12.39 | 0.00 | 5.13 | 0.37 |
| merge_samples_per_chrom | 23 | 0:00:51 | 47.64 | 509.40 | 38.99 | 42.01 | 19.36 | 49.58 | 66.33 | 36.76 |
| merge_samples_per_chrom_index | 23 | 0:00:03 | 18.66 | 1156.11 | 17.23 | 17.25 | 3.25 | 0.07 | 71.78 | 3.33 |
| rename_cram | 1 | < 0:00:01 | 0.00 | 0.00 | 0.00 | 0.00 | 0.00 | 0.00 | 0.00 | 0.00 |
| split_chroms | 115 | - | - | - | - | - | - | - | - | - |
| split_chroms_chr_string | 115 | - | - | - | - | - | - | - | - | - |
| tabbed_regions | 23 | - | - | - | - | - | - | - | - | - |
| tabbed_regions_chr_string | 23 | - | - | - | - | - | - | - | - | - |
| unique_regions_file | 23 | - | - | - | - | - | - | - | - | - |
| wget_cram | 1 | 1:19:52 | 4.48 | 9.36 | 4.48 | 4.48 | 3.94 | 14673.86 | 4.99 | 238.90 |

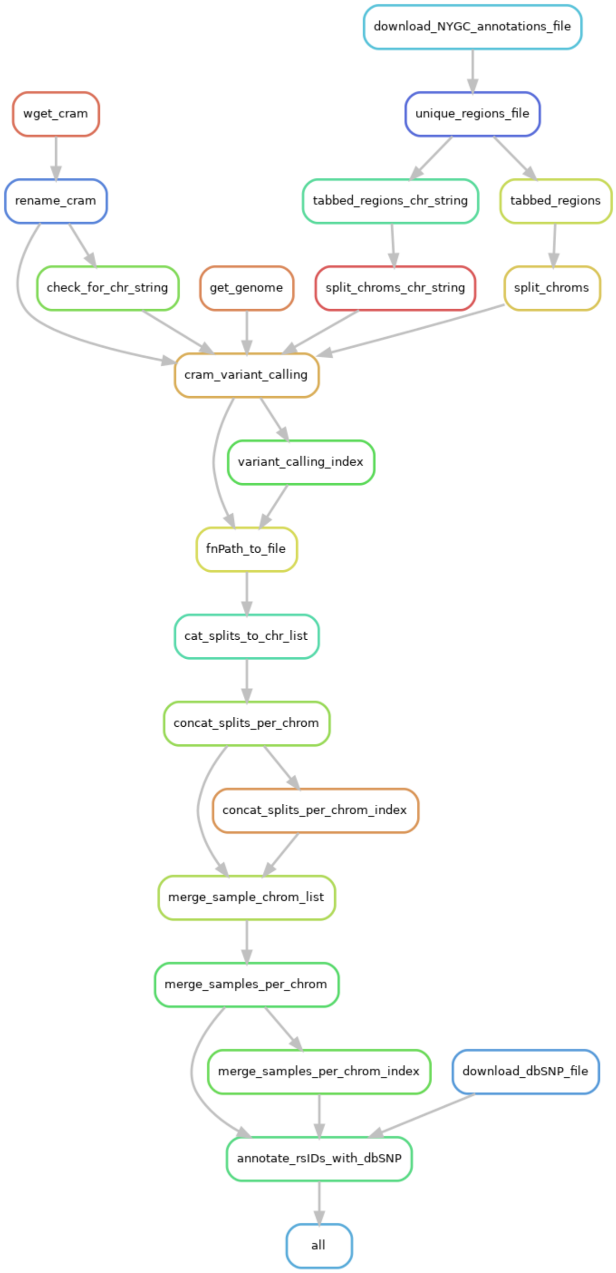


**Figure S2**. A directed acyclic graph for Iliad's stored sequence read data module.

**Table S3.** Time and resource usage for *Iliad*'s SNP Array data module. Each rule in the Snakemake workflow utilized the "benchmark" directive to collect this data. Some rules will constitute a number of jobs depending on the wildcards e.g. chromosomes and number of splits per chromosome. For this reason, values for time and resource usage were averaged across the jobs per rule. The benchmarks were conducted on three in-house IDAT samples on a dedicated remote server in the Department of Biology at Indiana University – Purdue University Indianapolis. This run was conducted using 8 cores provided to Snakemake for 8 jobs in parallel. No internal rule parallelization occurred. Since the "benchmark" directive is rule-specific for resource monitoring, there was no impact on any independent rule's time and resource report from performing these benchmarks with 8 jobs in parallel.

| Rule | Jobs | Average values across Jobs per Rule | | | | | | | | |
|---|---|---|---|---|---|---|---|---|---|---|
| | | h:m:s | max_rss | max_vms | max_uss | max_pss | io_in | io_out | mean_load | cpu_time |
| bwa_index | 1 | - | - | - | - | - | - | - | - | - |
| cleanRegionsFile | 1 | 0:00:22 | 2.14 | 45.77 | 1.65 | 1.69 | 0 | 0 | 0 | 0 |
| download_MEGA_physical_genetic_coordinates | 1 | 0:01:03 | 27.53 | 1571.78 | 27.06 | 27.08 | 0.07 | 0 | 0 | 0.03 |
| download_dbSNP_file | 1 | - | - | - | - | - | - | - | - | - |
| download_iaap | 1 | 0:00:15 | 0.02 | 1.22 | 0.03 | 0.03 | 0 | 0 | 0 | 0 |
| download_lociNames_conversion_file | 1 | 0:01:02 | 20.06 | 730.39 | 15.18 | 15.25 | 0 | 0 | 0 | 0.02 |
| download_product_files | 1 | 0:02:23 | 7.75 | 45.77 | 7.25 | 7.32 | 0 | 0 | 0 | 0 |
| extract_clean_regions | 1 | 0:10:44 | 28.54 | 755.38 | 27.9 | 27.93 | 0 | 0 | 0 | 0.02 |
| filter_for_dbSNP_vars | 1 | 0:21:10 | 0.02 | 1.22 | 0.03 | 0.03 | 0 | 0 | 0 | 0 |
| gencall | 1 | 0:01:34 | 8.27 | 45.77 | 7.86 | 7.93 | 0 | 0 | 0 | 0.01 |
| GenTrain_ClusterSep_QC | 1 | 0:00:12 | 21.78 | 742.04 | 21.13 | 21.19 | 0 | 0 | 0 | 0.01 |
| get_genome | 1 | - | - | - | - | - | - | - | - | - |
| gtc2vcf | 1 | 0:03:50 | 6.53 | 45.77 | 6.16 | 6.21 | 0 | 0 | 0 | 0.01 |
| insert_rsIDS | 1 | 0:00:18 | 6.7 | 45.77 | 0.02 | 0.02 | 0 | 0 | 0 | 0 |
| original_lociNames | 1 | 0:00:03 | 0.02 | 1.22 | 0.03 | 0.03 | 0 | 0 | 0 | 0 |
| replace_lociNames_to_rsIDs | 1 | 0:00:39 | 9.32 | 45.77 | 8.88 | 8.95 | 0 | 0 | 0 | 0.01 |

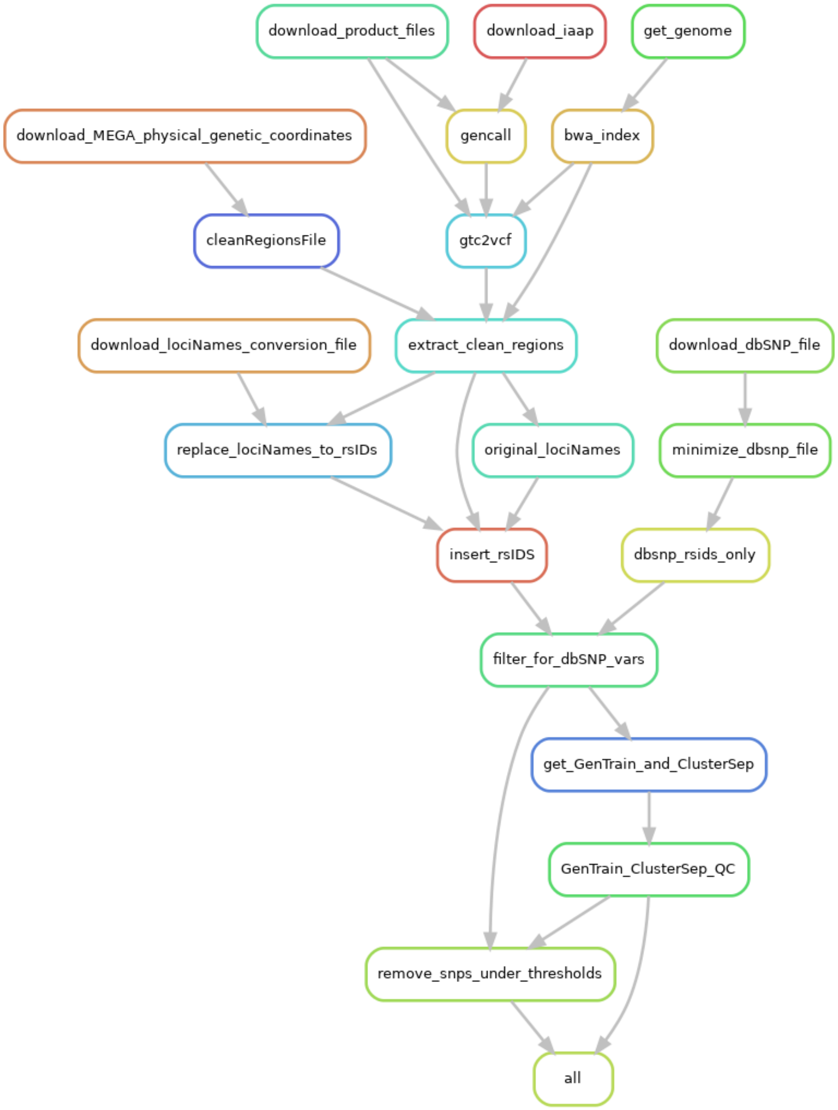


**Figure S3**. A directed acyclic graph for Iliad's SNP Array data module.

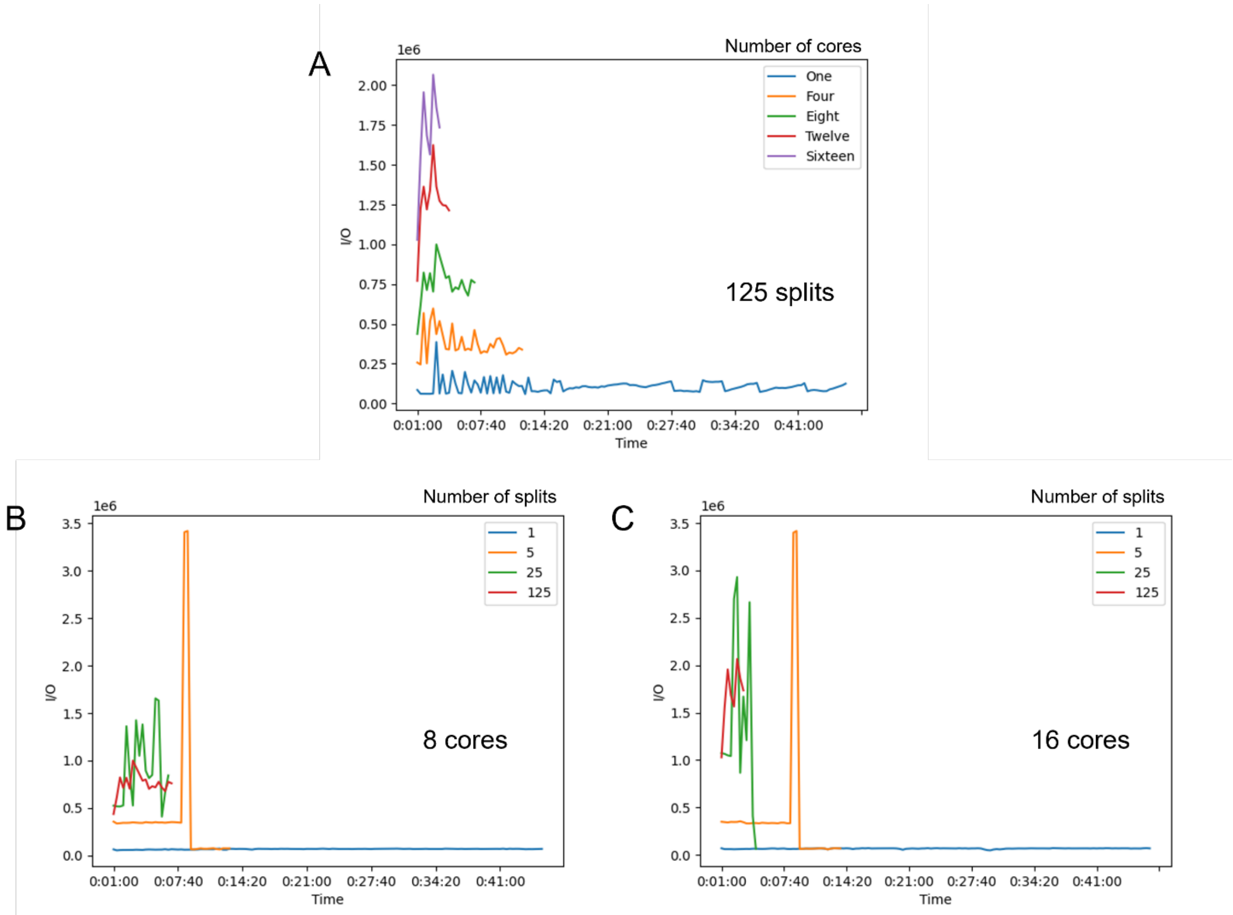


**Figure S4**. Time (h:m:s) and I/O (write data in KB) metrics for variant calling of one sample on chromosome 22 at **A)** one, four, eight, twelve, and sixteen cores to validate the logic that a large numbers of splits per chromosome, e.g. 125, performed faster with more supplied cores. Performance metrics for a varying number of splits at **B)** eight and **C)** sixteen cores to depict the lack of advantage of high splits with a lower number cores.
